# Integrated computational and experimental identification of *p53, KRAS* and *VHL* mutant selection associated with CRISPR-Cas9 editing

**DOI:** 10.1101/407767

**Authors:** Sanju Sinha, Karina Barbosa Guerra, Kuoyuan Cheng, Mark DM Leiserson, David M Wilson, Bríd M. Ryan, Ze’ev A. Ronai, Joo Sang Lee, Aniruddha J. Deshpande, Eytan Ruppin

## Abstract

Recent studies have reported that CRISPR-Cas9 gene editing induces a *p53*-dependent DNA damage response in primary cells, which may select for cells with oncogenic *p53* mutations^11,12^. It is unclear whether these CRISPR-induced changes are applicable to different cell types, and whether CRISPR gene editing may select for other oncogenic mutations. Addressing these questions, we analyzed genome-wide CRISPR and RNAi screens to systematically chart the mutation selection potential of CRISPR knockouts across the whole exome. Our analysis suggests that CRISPR gene editing can select for mutants of *KRAS* and *VHL*, at a level comparable to that reported for *p53*. These predictions were further validated in a genome-wide manner by analyzing independent CRISPR screens and patients’ tumor data. Finally, we performed a new set of pooled and arrayed CRISPR screens to evaluate the competition between CRISPR-edited isogenic *p53* WT and mutant cell lines, which further validated our predictions. In summary, our study systematically charts and points to the potential selection of specific cancer driver mutations during CRISPR-Cas9 gene editing.

---

CRISPR-Cas9 enables targeted gene disruption and editing, a powerful technology that expands our understanding of fundamental biological processes^1^. Beyond its impact on biological research, CRISPR-based approaches have been considered for various applications in medicine, from reparative editing of primary cells to the development of new strategies for treating a variety of genetic diseases, including cancer. However, several clinical trials based on CRISPR technology have been deferred due to significant potential risks, including off-target effects^2,3,4^, generation of unexpected chromosomal alterations^5^ and potential immunogenicity^6^. Other studies have demonstrated that double stranded breaks (DSBs) induced by CRISPR-Cas9 during gene knockout (KO) can lead to DNA damage response, whose level is associated with the copy number of the targeted gene.^7–10^.

Recent studies have shown that the DNA damage response following gene knockouts by CRISPR-Cas9 (CRISPR-KO) is mediated via *p53*, a known tumor-suppressor gene mutated in over 50% of cancers^11,12^. Haapaniemi *et al.* performed genome-wide CRISPR screening in immortalized human retinal pigment epithelial (RPE1) cells^12^, finding that a *p53*-mediated DNA damage response, followed by cell cycle arrest, is induced upon generation of DSBs by the Cas9 endonuclease, driving the selection of cells that have inactivated the *p53* pathway. Ihry *et al.* reported that the CRISPR-KO of even a single gene can induce a DNA damage response via the *p53* pathway^11^, with *p21/CDKN1A*, a canonical *p53* transcriptional target, playing a key role. These studies indicate that CRISPR-Cas9 genome editing techniques can potentially select for *p53* mutated cells pointing to an unanticipated oncogenic risk^11,12^. Since these studies were confined to a small number of primary or stem cell lines, a broader and more comprehensive evaluation of the potential mutation selection associated with CRISPR-based technology is warranted.

To address this challenge, we analyzed two large-scale genome-wide gene essentiality screens: *AVANA* (with CRISPR-Cas9)^10^, and *Achilles* (with shRNA)^13^. These screens were performed in a large panel of cancer cell lines from different tissues of origin and with a variable *p53* status^10,13^. We searched for individual genes whose silencing gives rise to significantly higher viability (reduced essentiality) in *p53* mutated vs *p53* wild-type (WT) cell lines, as the knockout (KO) of such genes can potentially lead to the selection of *p53* mutated cells. Importantly, we searched for genes that showed this effect only in the CRISPR-KO screens and not in the shRNA gene knockdown (KD) screens. The shRNA screens thus serve as a control enabling us to identify genes whose knockdown induces a bias towards *p53* mutated cells specifically upon CRISPR editing.

Our analysis of the CRISPR essentiality data was carried out on 248 cell-lines that are shared in both CRISPR and shRNA screens in DepMap^13^ (**Table S1**). We find 981 genes whose KO results in significantly higher cell viability in *p53* mutated (N=173) vs *p53* WT cell lines (N=75), while only 237 genes show the opposite trend (i.e. with significantly higher post-KO viability in *p53* WT cell lines, Methods). In contrast, the respective gene numbers related to *p53* status in the shRNA screens were balanced (∼1500 each, chi-squared test P<1.4E-284; the results in both cases are controlled for gene copy number as a potential confounding factor, Methods, **Figure 1a** top panel, **Figure S1**). Since the contribution of some of these mutations to p53 function has not been established, we repeated the above analysis focusing on cell lines harboring known loss-of-function *p53* mutations solely (Methods, N=78). Comparing those with the WT cell-lines as controls (N=75), we observed an even higher significance in the differences of median post-KO/KD cell viability (chi-squared test P=1.7E-292, **Table S2**, see Methods). These findings were further corroborated by a permutation test where the cell line’s *p53* mutation status was shuffled 10,000 times (P<1E-4, Methods).

**Figure 1.**
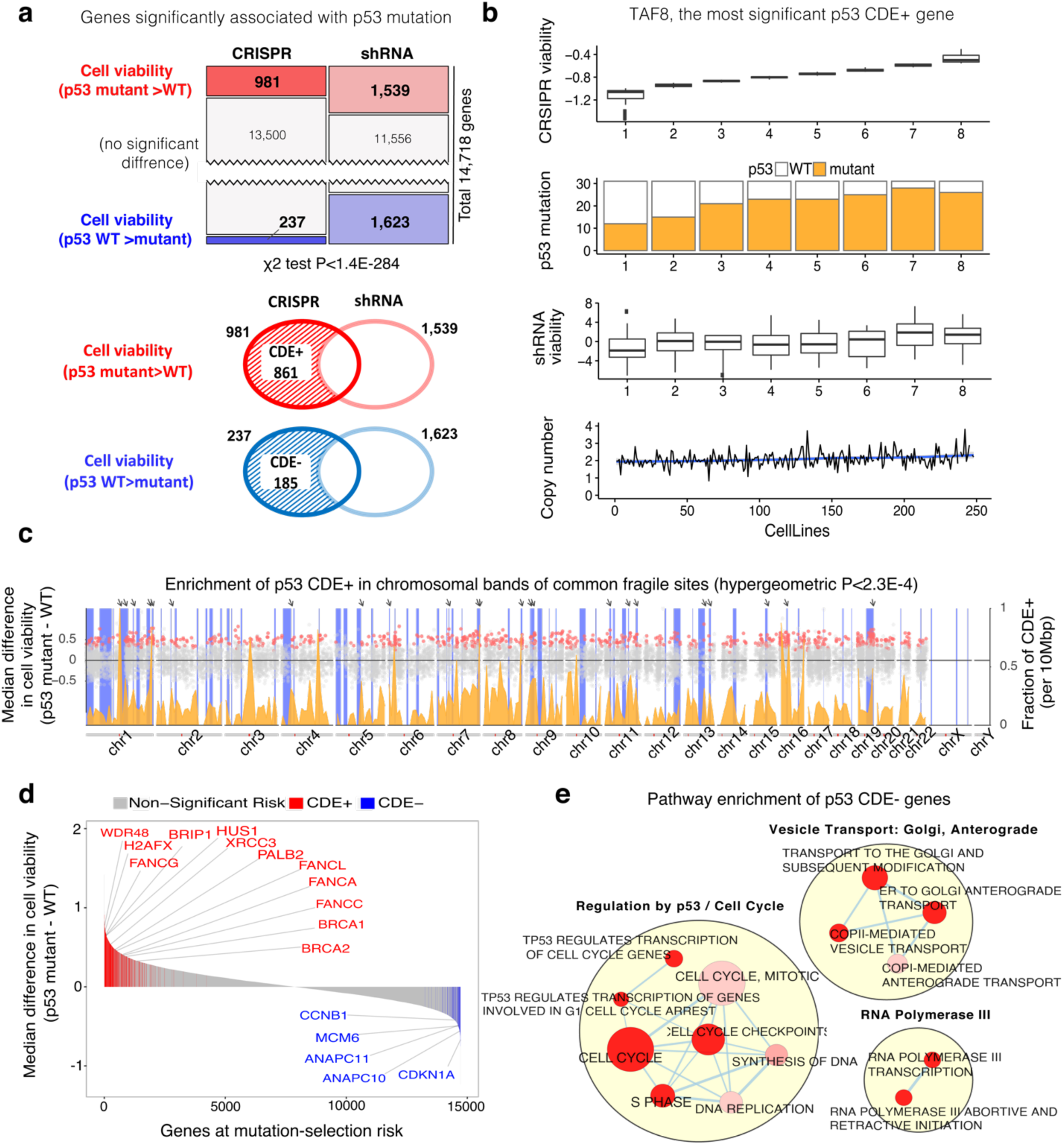
A genome wide view of *p53*-associated oncogenic risk. **(a)** Upper panel: number of genes whose essentiality is significantly associated with *p53* mutation status in CRISPR and shRNA screens (one sided Wilcoxon rank-sum has been performed with FDR threshold of 0.1). Lower panel: the definition of CDE+ and CDE− genes. **(b)** The distribution of *TAF8* (the top CDE+ gene of *p53*) post KO/KD viability scores in the CRISPR (top panel) screen as a function of *p53* mutation status. The cell lines were divided into 8 bins of equal size, which are ordered by their viability after the CRISPR-KO of *TAF8*. The fraction of *p53* mutant cell lines in each bin is plotted on the second panel. The distribution of *TAF8* post shRNA-KD viability scores is displayed on the third panel as a control. The fourth, lowest panel visualizes that the copy number of *TAF8* is about the same in the different bins. **(c)** Enrichment of *p53* CDE+ genes in common fragile sites (CFSs). The x-axis denotes the chromosomal position; the scatter plot (y-axis on the left-hand side) shows the difference of median post-CRISPR-KO cell viability values in *p53* mutant vs *p53* WT cell lines for *p53* CDE+ genes (red dots) and all other genes (grey dots); the density plot (colored orange, y-axis on the right-hand side) shows the fraction of *p53* CDE+ genes among all genes per DNA segments of 10 Mbp along the genome; the vertical blue bars indicate the chromosomal bands of CFSs, and prominent sites where peaks of high CDE+ gene density coincide with CFSs are marked by arrows on the top. **(d)** The distribution of predicted CRISPR-risks across the genome. Significant CDE+ genes that are part of FA pathway are marked in red and significant CDE− genes that are part of cell cycle regulation in blue. **(e)** Visualization of the pathways enriched for *p53* CDE− genes. Only significantly enriched pathways (FDR<0.1) specific to CRISPR (and not in genes showing differential essentiality in the shRNA screens are shown). Pathways are depicted as nodes whose sizes correlate with pathway lengths and colors represent enrichment significance (the darker, the more significant). Pathway nodes are connected and clustered based on their functional similarities, and higher-level functional terms are given for each of the clusters (Methods). For clarity, only the largest clusters are shown.

Among the 981 genes whose CRISPR-KO results in higher cell viability when *p53* is mutated, a vast majority (87%, 861 genes) show this differential essentiality effect only in CRISPR screens and not in shRNA screens, termed *CRISPR-specific differentially essential positive (CDE+)* genes (illustrated in the lower panel of **Figure 1a;** CDE+ genes listed in **Table S3A**; this result holds when controlling for copy number, **Figure S1**). We show the distribution of cell-lines’ *p53* status and copy number with the cell-lines ordered by cell viability after CRISPR-KO of the top CDE+ gene, *TAF8*, as an example (**Figure 1b**). We then investigated whether there was a selective enrichment for functional or architectural features in the CDE+ sgRNAs and their cognate genes. Notably, analysis of genomic localization of sgRNAs demonstrates that CDE+ genes are preferentially located in chromosomal bands containing common fragile sites (CFSs) (hypergeometric P<2.3E-4, **Figure 1c, Table S4**). These chromosomal regions are prone to replicative stress, fork collapse and DNA breaks that cause genomic instability^14^ (not observed for the genes showing differential essentiality only in the shRNA screens). As CRISPR-KO events can induce DNA damage, including kilobase-scale structural alterations near the targeted site^15^, this finding supports the possibility that the formation of a CRISPR-induced DSBs in the vicinity of CFS bands may promote the activation of a *p53*-dependent cell death response. Reduced cell fitness may then result in the selection of *p53* mutated cells where this response is blunted. Secondly, we find that CDE+ genes are also enriched for genes whose sgRNAs target highly accessible chromatin (methods; hypergeometric P<0.02), consistent with the findings of a recent work that double-stranded breaks induced in highly accessible chromatin regions could induce stronger DNA damage response^16^.

Next, our analysis shows that CDE+ genes are significantly enriched for factors associated with DNA damage response (FDR-corrected hypergeometric P<0.01, **Table S3B**). Some of the most significant CDE+ genes are involved in DNA repair such as BRCA1, BRCA2 as well as Fanconi anemia (FA) core complex genes FANCG, FANCG and FANCC. This is consistent with the notion that inactivating these DNA repair genes enhances DNA damage induced by Cas9, promoting cell death in *p53* wild-type cells. It further suggests that the recently reported involvement of the FA pathway in the repair of DSBs induced during CRISPR gene-editing^17^ may have a *p53* dependency. **Figure 1d** shows the median differences of post-KO viability between *p53* mutant vs *p53* WT cell lines for each CDE+ gene, termed *CRISPR-KO risk*. The top hits are marked, including genes located in CFSs and those involved in DNA repair (Methods). As evident, the magnitude of the differential viability scores observed for top ranked CDE+ genes is high, suggesting that a *p53* mutated cell population is indeed quite likely to be selected for.

In addition to the CDE+ genes, we identified a group of 237 genes that are more essential in CRISPR screens in *p53* mutated than WT cell lines. Importantly, most of these (78%; 185 genes) show this pattern of *p53* mutated differential essentially only in the CRISPR screen and not in shRNA screens, thus termed CRISPR-specific differentially essential negative *(CDE−)* genes (illustrated in the lower panel of **Figure 1a**). Consistent with earlier reports^11,12^, the top ranked CDE− gene is *CDKN1A* (a.k.a. *p21*, Wilcoxon rank sum P<1.85E-08, **Figure 1d, Table S3A**), a *p53*-controlled cyclin-dependent kinase inhibitor implicated in the control of cell cycle progression. Generally, CDE− genes are involved in cellular processes that engage *p53*, including mitotic checkpoints, DNA replication and cell cycle (**Table S3B**, visualized in **Figure 1e**, FDR corrected hypergeometric P<0.1). Transiently inhibiting CDE− genes concomitantly with the CRISPR editing of other desired target genes may mitigate the mutation selection of the latter, and hence could be of potential interest from a translational point of view.

Next, we experimentally tested whether CDE+ and CDE− genes show preferential selection for mutant and WT *p53* cells, respectively, in an isogenic experimental setting. An isogenic *p53* wildtype and *p53* mutant pair of the MOLM13 leukemia cell line were established for pooled CRISPR-KO screening using a CRISPR library targeting the top *p53* CDE+ and CDE− genes. To ensure that the CDE selection effects were specific to CRISPR-KO, the same genes were also targeted by a pooled CRISPR-interference (CRISPRi) library using a catalytically inactive Cas9, fused to the KRAB repressor and the methyl CpG binding protein MeCP2^18^ (illustrated in **Figure 2a**). Since our initial comparison of CDE essentiality was conducted between the CRISPR-KO and shRNA screens, we set to eliminate inherent differences between CRISPR-Cas9 and shRNA-based approaches by performing the CRISPR-KO and CRISPRi of the same genes with equal number of sgRNAs in an isogenic setting. From this screen, we identified the genes showing higher essentiality in *p53*-WT vs *p53* mutant cells, but only in CRISPR-KO and not CRISPRi. We confirmed that these genes are enriched for the *p53* CDE+ genes identified earlier in the overall analysis (hypergeometric P<1E-8, Methods). A parallel enrichment of the predicted CDE− genes was confirmed using a similar approach (hypergeometric P<2E-4). The CDE+ genes are differentially more essential in wildtype than mutant, specifically in CRISPR-KO but not in CRISPRi (Wilcoxon P<1E-06 for CRISPR-KO, P<0.32 for CRISPRi). The effect size for the top 10% of CDE+/− genes identified earlier is depicted in **Figure 2b** (Methods).

**Figure 2.**
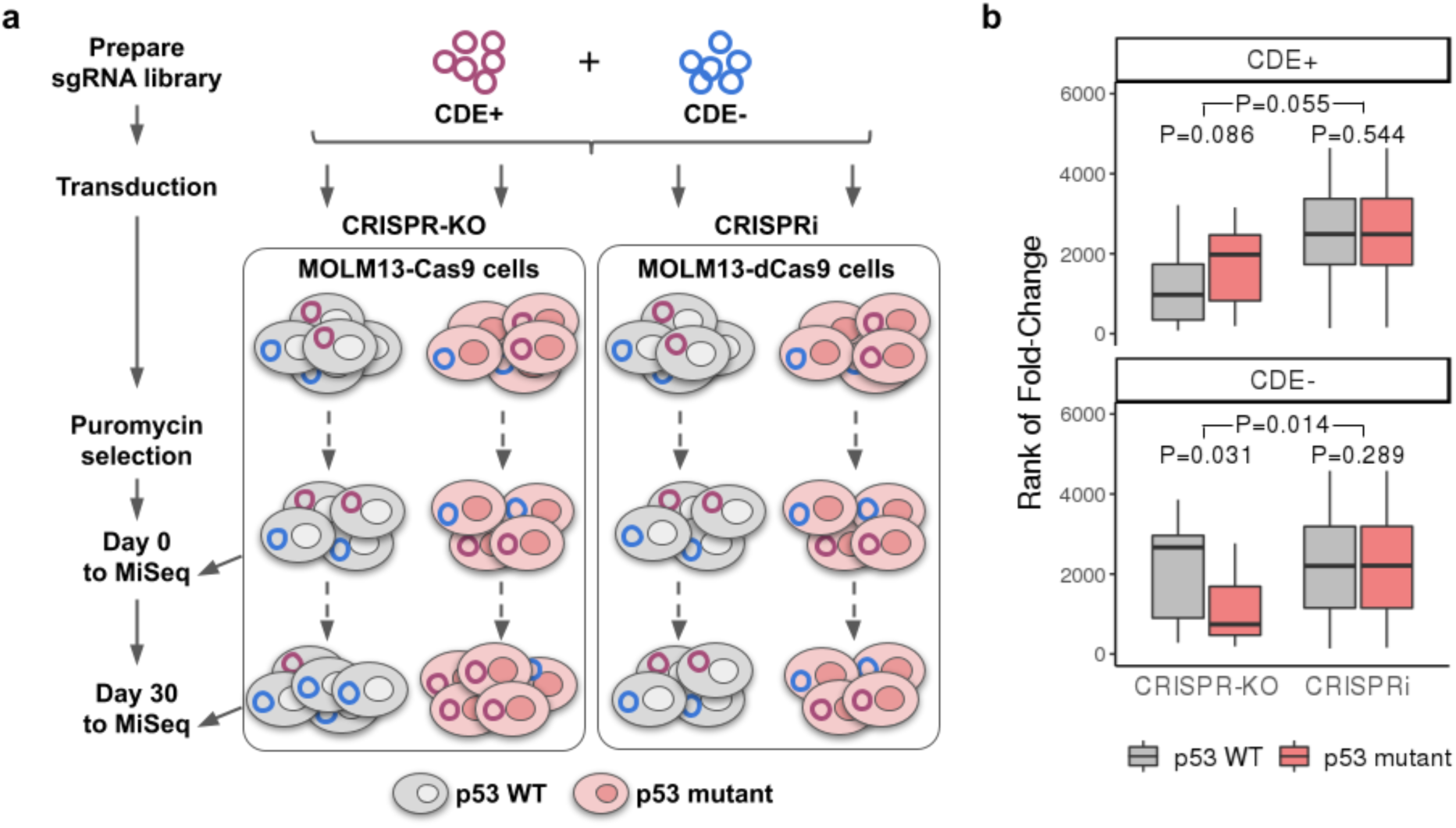
Validation of *p53* CDE genes in isogenic MOLM13 cell lines via pooled CRISPR screens. **(a)** A flowchart showing the experimental procedure of CRISPR-KO and CRISPRi screening of pooled *p53* CDE+/− genes in a pair of *p53*-isogenic MOLM13 cell lines. See Methods for details. **(b)** The day 30 to day 0 fold-change (converted to rank) of reads corresponding to the sgRNAs for *p53* CDE+ genes (upper panel) and CDE− genes (lower panel), in *p53* WT MOLM13 cells (gray boxes) vs the isogenic *p53* mutant cells (red boxes) for the CRISPR-KO and CRISPRi screenings, respectively. The bottom P values are for Wilcoxon signed-rank tests comparing *p53* WT and mutant cells, the upper ones are P values of non-parametric tests comparing the difference of *p53* mutant and WT rank values between CRISPR-KO and CRISPRi experiments.

We then tested whether the CRISPR-KO of CDE+ genes leads to selection of *p53* mutant cells in a competitive setting. Towards this end, the top 5 predicted CDE+ genes were knocked-out using distinct sgRNAs in wild-type and isogenic MOLM13-*p53* cells. Following a lentiviral sgRNA transduction the wild-type and mutant cells were mixed at an initial ratio of 95:5, respectively and monitored by flow cytometry up to 25 days for the proportion of *p53* wild-type (TdTomato-) and *p53* mutant (TdTomato+) cells (experimental design illustrated in **Figure 3a**). Two out of the five CDE+ genes (NDUFB6 and NDUFB10) exhibited a significant and strong *p53* mutant selection, across several independent sgRNAs, whereby the *p53* mutant population was progressively enriched, up to five-folds over the *p53* wild-type cells at day 25 (**Figure 3b**, blue lines). Increase of *p53* mutant cells was not observed using non-targeting control (NTC) sgRNA (**Figure 3b**) and no inverse enrichment in *p53* wild-type cells was observed in the competitive essays involving the three other CDE+ genes (**Figure S2**). Importantly, a parallel competitive assay for knock down of these CDE+ genes with CRISPRi did not result in outgrowth of *p53* mutant cells (**Figure 3b**, pink lines). Notably, the initial abundance of *p53* mutated cells used in our competitive assays is comparable to the baseline *p53* mutation rate (3.5%) seen in human pluripotent stem cells (hPSCs) typically used in CRISPR-Cas9 based therapeutic editing^19^.

**Figure 3.**
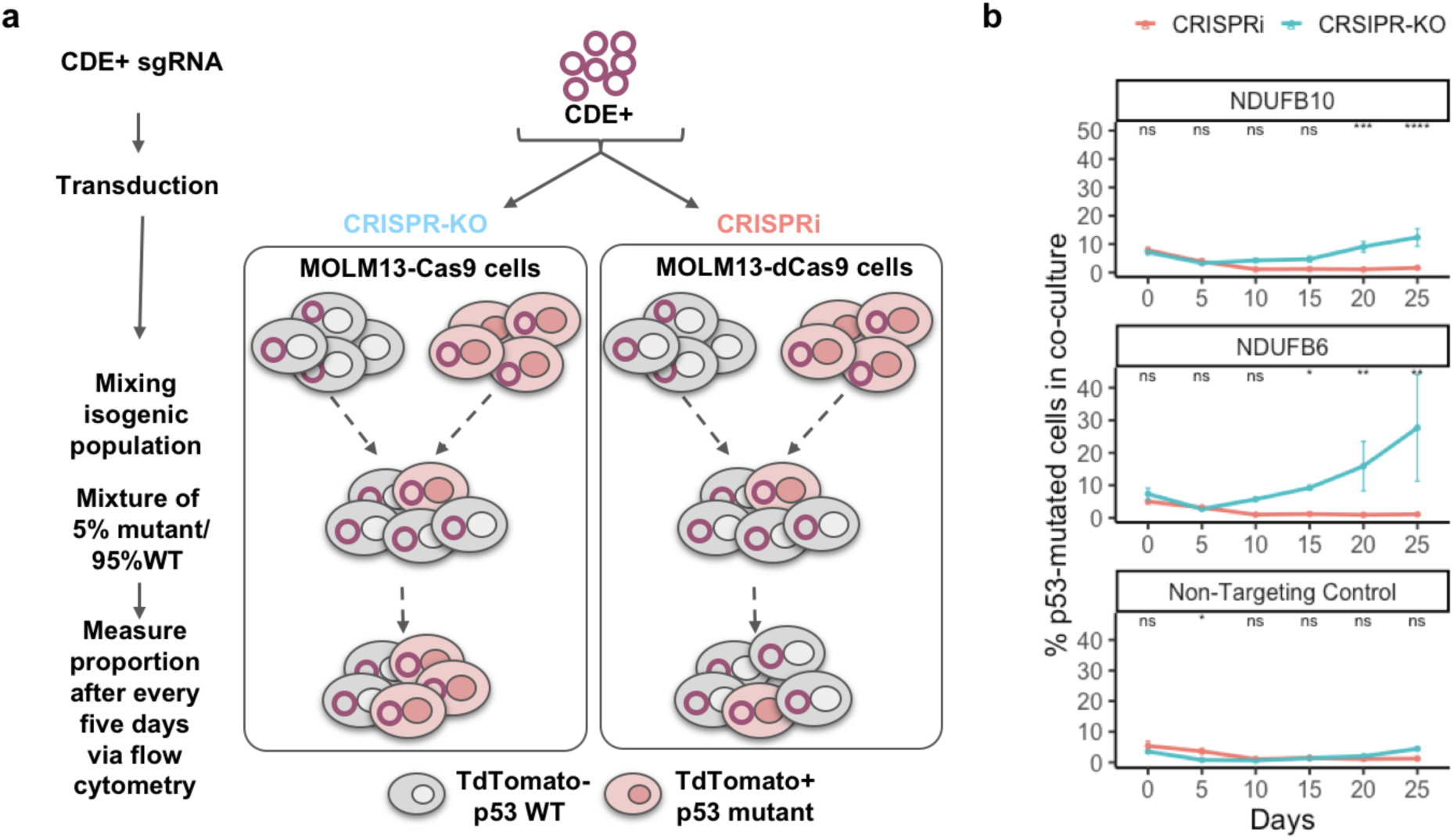
Selection for *p53* mutant cells under CRISPR-Cas9 knockout of CDE+ genes in a co-culture of *p53* WT/mutant cells. **(a)** An illustration showing the experimental design of the competition assay where isogenic *p53* WT/mutant MOLM13 cell lines were mixed with a ratio of 5:95 and top *p53* CDE+ genes were knocked out by CRISPR-Cas9. Population ratio was monitored for 25 days at a five-day interval starting from the day of sgRNA transduction. **(b)** Change of ratio of *p53* mutant and WT cells (Y-axis, % *p53* Mutant/WT Cells) with time (number of days on the X-axis), where days are color coded. The leftmost panel (NTC) shows the corresponding result in the control experiment using non-targeting sgRNAs.

To further assess whether similar effects can be observed in primary cells, we analyzed the original genome-wide screens conducted by Haapaniemi *et al.*^*12*^ in the setting of *p53* isogenic non-transformed RPE1 cells. We identified the genes that are more (or less) essential in WT vs isogenic *p53* mutant cells (termed DE+ and DE-genes respectively, Supp. Note 1). Consistent with our findings the top 100 DE+/− genes were significantly enriched for our list of *p53* CDE+/− genes respectively (hypergeometric test P<0.04 for both CDE+/−), where this significance robustly holds true (P<0.1) for a wide range of top DE+/− and CDE+/− genes (Supp. Note 1). These results testify that our CDE+/− genes findings from the analysis of CRISPR KO in cancer cell lines coincide in a significant manner with those found in primary human cells.

Given that the CRISPR-KO of CDE+ genes preferentially reduces the viability of *p53* WT cells, we hypothesized that copy number alterations in CDE+ genes (which could act as a possible surrogate for number of DSBs) could also reduce the fitness of *p53* WT tumors. To test this hypothesis, we analyzed the somatic copy number alteration (SCNA) and patient survival data of 7,547 samples from the cancer genome atlas (TCGA)^20^. As a control, we used genes whose essentiality is not associated with *p53* mutational status (Methods). We find that the SCNA (both amplifications and deletions, Methods) in CDE+ genes is significantly lower in *p53* WT tumors compared to that of control genes (Wilcoxon rank sum P<2.7E-5). This suggests that copy number variations in CDE+ genes are selected against specifically in *p53* WT tumors, as *p53*-mediated responses are detrimental to their fitness. Notably, this was not the case for *p53* mutant tumors or for CDE− genes (Wilcoxon rank sum P>0.5). In addition, we observe that the magnitude of SCNA events of CDE+ genes is associated with enhanced patient survival in *p53* WT but not in *p53* mutated tumors (hypergeometric P<0.046, Methods), further supporting the notion that these amplification/deletion events reduce the fitness of these *p53* WT tumors. Taken together, these observations provide further evidence from patients’ tumors for the functional risks incurred by the inactivation of CDE+ genes.

While recent studies pointed to the oncogenic risk involving *p53*^*11,12*^, an intriguing open question is whether CRISPR knockouts may also select for mutations in other major cancer driver genes. To identify such potential additional cancer drivers, we applied the analysis described above for p53 to each gene in the list of cancer driver genes provided by Vogelstein *et al.*^*21*^ (focusing on those genes that are mutated in at least 10 of the screened cell lines, N=61). We identified the oncogene *KRAS* and the tumor suppressor *VHL* (Von Hippel-Lindau) as such *master CDE regulators* (or “master regulators” for short, Methods, **Figure 4a, Table S5**). That is, like *p53*, each of these two genes have a CDE+/− skewed distribution in the CRISPR compared to the shRNA screens. The CDE+ and CDE− genes of each of these master regulators are listed in **Table S3A**. Some genes are classified as CDE+ of multiple master regulators, and thus their CRISPR-KO may impose considerable risk of selecting for more than one mutated cancer gene. E.g., *SREK1* is both a *p53* and *VHL* CDE+ gene; fourteen genes are CDE+ genes of both *KRAS* and *p53*; and ten genes are CDE+ genes of both the *KRAS* and *VHL* (**Table S6**).

**Figure 4.**
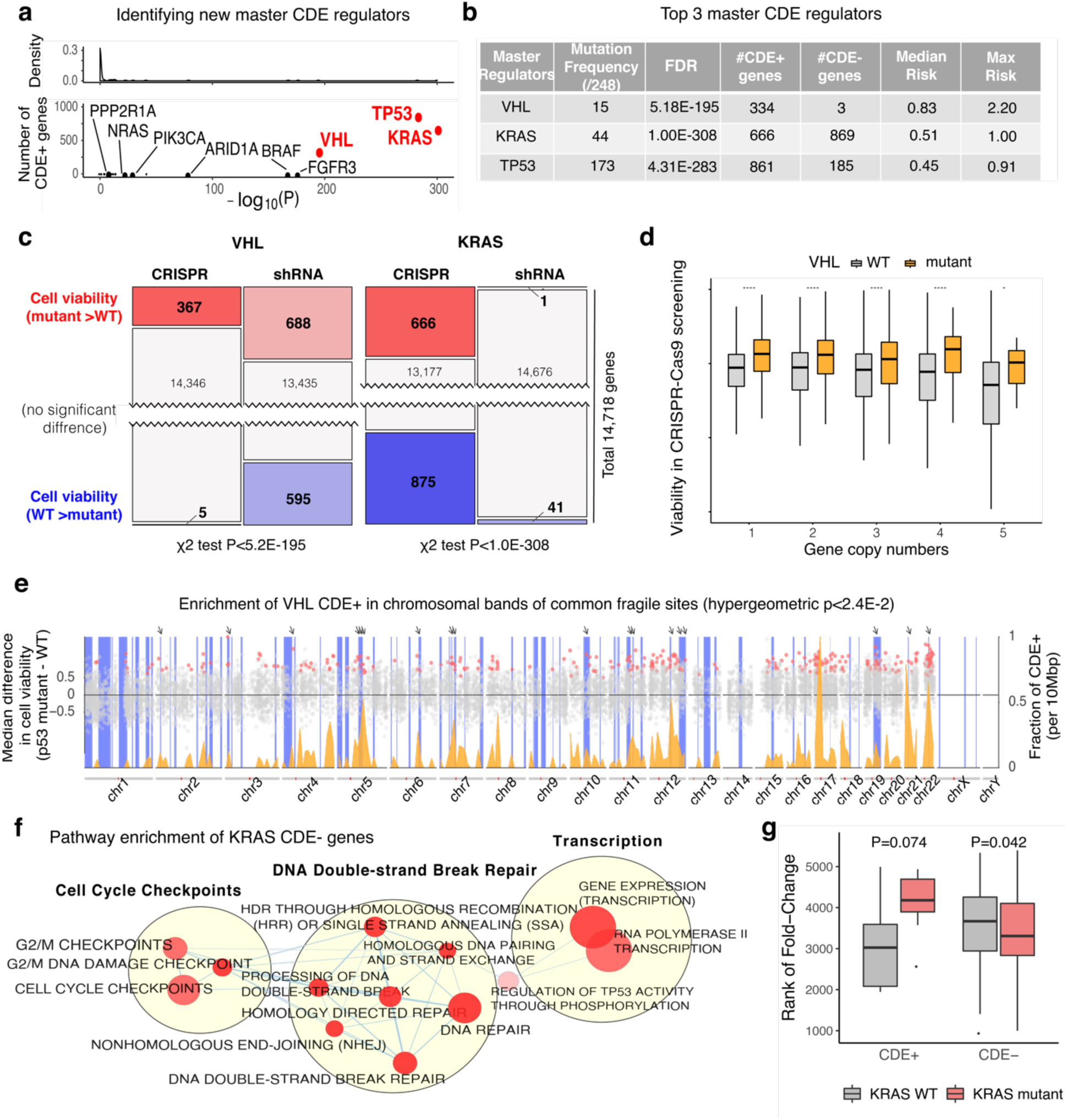
Beyond *p53*: *VHL* and *KRAS* as master CDE regulators. **(a)** Top panel: A density plot showing that almost all cancer drivers are not identified as significant master CDE regulators. The x-axis denotes their significance (negative log-transformed P value of the chi-squared test for being a master regulator, see Methods). Bottom panel: Displays the identity of the nine genes having significant P values and more than one CDE+ gene; only the top three most significant genes (marked in red) have a considerable number of CDE+ genes, and hence we have focused on studying their role as master regulators further. **(b)** A table describing key properties of the three master regulators. The first column denotes their names, the second column displays the frequency of their non-synonymous mutations among 248 cell lines, the third column displays the FDR corrected Fisher’s exact P value denoting the significance of the CRISPR-specificity of the association of gene essentiality with their mutation status. The fourth and fifth columns denote the number of CDE+ and CDE− genes identified for each of the master regulators. The sixth and seventh columns denote the median and maximal risk score (difference in cell viability after CRISPR-KO of the CDE+ genes between the mutant and WT cell lines of the respective master regulators). **(c)** The number of genes whose essentiality is significantly associated with *VHL* (left) and *KRAS* (right) mutational status in CRISPR and shRNA screens (one sided Wilcoxon rank-sum has been performed with FDR threshold of 0.1). **(d)** The effect of *VHL* mutation status on CRISPR-KO viability (y-axis) is significant across a wide range of CDE+’s copy numbers (x-axis). CN=1 denotes cases with copy number less than or equal to 1 and CN=5 denotes cases with copy number greater than or equal to 5. The number of stars at the top of boxplot represents the significance of the difference (one-sided Wilcoxon rank sum test). *P<0.05, **P<0.01, ***P<0.001, ****P<0.0001. **(e)** The enrichment of *VHL* CDE+ genes in common fragile sites (CFSs). The x-axis denotes the chromosomal position; the scatter plot (y-axis on the left-hand side) shows the difference of median post-CRISPR-KO cell viability values in *VHL* mutant vs WT cell lines for *VHL* CDE+ genes (red dots) and all other genes (grey dots); the density plot (colored orange, y-axis on the right-hand side) shows the fraction of *VHL* CDE+ genes among all genes per DNA segments of 10 Mbp along the genome; the vertical blue bars indicate the chromosomal bands of CFSs, and prominent sites where peaks of high CDE+ gene density coincide with CFSs are marked by arrows on the top. **(f)** Visualization of pathways enriched for *KRAS* CDE− genes. Only significant pathways (FDR<0.1) specific to CDE and not to the genes showing differential essentiality in the shRNA screens are included. Pathways are shown as nodes whose sizes correlate with pathway lengths and colors represent the significance of their enrichment (the darker the more significant). Pathway nodes are connected and clustered based on their functional similarities, and higher-level functional terms are given for each of the clusters (Methods). For clarity, only the largest clusters are shown. **(g)** CRISPR-Cas9 screens of the top *KRAS* CDE gene knockouts were performed in isogenic MOLM13 and MOLM13-*KRAS*-G12D cell lines. The box plot shows that the sgRNAs of the *KRAS* CDE+ genes become significantly more depleted in *KRAS* WT cells vs *KRAS* mutant cells, vice versa for *KRAS* CDE− genes. The P value of one-tailed Wilcoxon signed-rank test is shown.

As evident from **Figure 4b**, the two additional master regulators uncovered by our analysis have considerably higher predicted median mutant selection levels than *p53;* that is, the CRISPR-KO of their CDE+ genes is likely to drive higher levels of mutant selection than KO of the CDE+ genes of *p53*. Notably, for *VHL*, although the absolute number of its CDE+ genes is smaller than that of *p53*, the ratio of its CDE+ to CDE− genes is strikingly higher than that observed for *p53*, while the parallel distribution observed in the shRNA-KD screens is balanced (**Figure 4c**). Just like *p53*, the mutational status of *VHL* is significantly associated with the essentiality of CDE+ genes independent of copy number (**Figure 4d**). The CDE+ genes of *VHL* are also enriched in chromosomal bands of CFSs (**Figure 4e and Table S4**, hypergeometric P<2.4e-2). Indeed, *VHL* can act as a positive regulator of *p53* in DNA damage-induced cell cycle arrest or apoptosis^22^, possibly accounting for its role as a master regulator.

For *KRAS*, a major oncogene that is involved in cell signaling, we find high numbers of CDE genes, while only very few *KRAS* mutation-associated genes are identified in the shRNA-KD screens (**Figure 4c**). Mutated *KRAS* is known to activate various DNA repair pathways and may override the trigger of cell death upon DNA damage^23,24^, supporting its role as a master regulator. Consistently, the CDE− genes of *KRAS* are significantly enriched for DNA DSB repair pathways (FDR<0.02, Methods, visualized in **Figure 4f, Table S3**). As observed for *VHL and p53* (**Figure 4d and S1**, respectively), the effects of *KRAS* are also independent of gene copy number (**Figure S1**). Consistent with these findings, mining the data from genome-wide CRISPR and shRNA screens performed in two *KRAS* isogenic cell lines (DLD1 and HCT116)^25^ revealed that the predicted *KRAS* CDE+/− genes have marginally significant overlap with the DE+/− genes identified in both the cell lines CRISPR screens (hypergeometric test P<0.1, Supp. Note 2) and not in the corresponding shRNA screens.

Next, similar to our experiments in the case of *p53*, we generated a focused sgRNA library consisting of 10 sgRNAs for each of the top 186 *KRAS* CDE+/− genes and performed CRIPSR-Cas9 screen in a pair of WT and isogenic *KRAS* G12D mutant MOLM13 cell lines (Methods). We confirmed that DE+/− genes derived from this experiment are highly enriched in our previously identified *KRAS* CDE+/− genes from pooled screens (Methods; hypergeometric P=0.002 for CDE+ and P=0.006 for CDE−). The *KRAS* CDE+ genes were found to be differentially more essential in wildtype than in mutant, as expected (Wilcoxon paired P=9E-04). The effect size exhibited by the top 50% CDE+/− genes is illustrated in **Figure 4g** (Methods).

In all, our study established three master regulators (MRs) and their CDE+ genes that may mediate the potential risk of CRISPR-Cas9-associated mutant selection. Based on our results there can be multiple factors may contribute to the identity of these CDE+ genes. First, some CDE+ genes are involved in DNA damage repair. Second, other genes are located within chromosomal common fragile sites (CFSs) or highly accessible chromatin regions (**Figure 1c & 4e**). Indeed, targeting such CDE+ genes is likely to result in more severe DNA damage and consequently preferentially kill cells with intact DNA repair processing. Overall, we find that these factors can together account for up to 15% of the CDE+ genes we have identified (i.e. proportion of CDE+ genes within CFS sites). Additionally, a CDE+ gene could potentially exhibit its differential lethality effects because the CRISPR-Cas9 sgRNAs which is targeting it have higher off-target effects, thus causing more DNA damage. To assess this possibility, we calculated an off-target score for each sgRNA (Supp. Note 3). We find that the CDE+ genes of *p53* and *VHL* (but not those of *KRAS*) are enriched for genes with high sgRNA off-target scores (hypergeometric P<6.9E-06 for *p53*, P<6.5E-02 for *VHL*; Methods; Supp. Note 3). This indicates that sgRNA off-target effects may indeed underlie some of the CDE+ genes we identified. Reassuringly though, we find that off-target effects can account for no more than 10% of the CDE+ genes (Supp. Note 3), ruling out the possibility that such technical limitations have markedly biased our results. Taken together, as these three putative mechanisms can explain about 25% of the CDE+ genes we have identified, the mechanisms underlying the rest are yet open to further studies.

Taken together, our findings raise several considerations of potential relevance for therapeutic gene-editing applications using the CRISPR-Cas9 system. Firstly, our studies confirm computationally and experimentally that the selection for *p53* mutations induced by CRISPR-Cas9 previously demonstrated in a primary human cell type^11,12^ also occurs in hundreds of cell lines from diverse tissues of origin. Second, our analysis indicates that genomic editing by CRISPR-Cas9 may also induce selective enrichment of cells with mutations in *KRAS* and *VHL* – central drivers of human malignancies. Our study indicates that selective outgrowth of mutated cells may be accelerated by CRISPR-KO of CDE+ genes. We validated this selection advantage of CDE genes in a more focused CRISPR screens of isogenic cell lines. Notably, using a competition assay, we directly demonstrate for the first time that the knockdown of top CDE+ genes indeed induces the selective growth advantage of *p53*-mutant cells. It is important to note that while our analysis is performed in cancer cell lines, the resulting CDE genes have a significant overlap with the findings in human non-cancerous cells reported earlier^11,12^. Notably, we find evidence that the CRISPR-related KO effects identified *in vitro* may have an echo that is traceable in gene copy number alterations observed *in vivo*, modulating the evolution of patients’ tumors. In closing, our results further expand upon and strengthen the previous findings^11,12^, suggesting that CRISPR-Cas9 editing or knockout of CDE+ genes may select for mutations in the cancer genes *p53, KRAS* and *VHL*.

## Methods

### CRISPR and shRNA essentiality screen data

We obtained CRISPR-Cas9 essentiality screen (or dependency profile) data in 436 cell lines from *Meyers et al.*^*10*^ for 16,368 genes, whose expression, CNV and mutation data are available via CCLE portal^26^. We obtained the shRNA essentiality screen data in 501 cell-lines from DepMap portal^27^ for 16,165 genes, whose expression, CNV and mutation data is available publicly via CCLE portal^26^. The 248 cell-lines and 14,718 genes that appear in both datasets were used in this analysis (**Table S1**). For mutation data, only non-synonymous mutations were considered. Synonymous (silent) mutations were removed from the pre-processed MAF files downloaded from CCLE portal^26^.

### Identifying CRISPR specific differentially essential genes of a master CDE regulator

For a given master CDE regulator (e.g. *p53*), we checked which gene’s essentiality (viability after knockout) is significantly associated with the mutational status of the master regulator using a Wilcoxon rank sum test in the CRISPR and shRNA datasets, respectively (FDR<0.1). *CRISPR-specific differentially essential (CDE) genes* denotes two sets of genes depending on the direction of the association with the mutation. CDE+ genes are those whose CRISPR-KO is significantly more viable when the master regulator is mutated while their shRNA silencing is not, whereas CDE− genes are those whose CRISPR-KO is significantly more viable when the master regulator is WT while their shRNA silencing is not. We filtered out any candidate CDE genes whose copy number was also significantly associated with the given mutation to control for potentially spurious associations coming from copy number (we removed genes showing significant association (FDR<0.1)) – the exact procedure used is described below in the section titled “*Identifying master CDE regulators of CRISPR-KO*”).

### Identifying CDEs associated with p53-loss-of-function mutations

Out of a total of 248 cell lines that we analyzed, 173 cell lines (69.7%) have *p53* non-synonymous mutations. In addition to identifying CDEs by considering all non-synonymous mutation, we additionally employed a more conservative approach where we aimed to consider only *p53 loss-of-function* (LOF) mutations in the CDE identification process. To this end, we considered a mutation to be LOF if it was classified as non-sense, indel, frameshift, or among the 4 most frequent non-functional hotspot mutations (R248Q, R273H, R248W and R175H within the DNA-binding domain, determined as pathogenic by COSMIC^28^). Using this definition we obtained new mutation profiles for *p53* and identified CDE genes via the same method described in the section titled “*Identifying CRISPR specific differentially essential genes* of a master CDE regulator.”

### Identifying master CDE regulators of CRISPR-KO

To identify additional master regulators like *p53*, we considered 121 cancer driver genes identified by Vogelstein *et al.*^*21*^, whose nonsynonymous mutation is observed in at least 10 cell lines (N=61). We determined whether each of these genes is a master CDE regulator as follows: for each of the 61 candidate genes, we tested the association between the essentiality of each of genes in the genome (reflected by post-KO cell viability) with the mutational status of the candidate master regulator gene using a Wilcoxon rank sum test. We then counted the number of genes, whose essentiality is: (i) significantly positively associated with the candidate master regulator mutational status (FDR-corrected p-value<0.1, median essentiality of WT>mutant of the cancer gene), (ii) significantly negatively associated with the candidate master regulator mutational status (FDR-corrected p-value<0.1, median essentiality of WT<mutant of the cancer gene), and (iii) not associated (FDR-corrected p-value>0.1) with the candidate master regulator mutation status; we performed this computation separately for the CRISPR and the shRNA screens, respectively. This computation results in a 3-by-2 contingency table for each candidate master regulator gene. We then checked whether the distribution of the above three counts in CRISPR dataset significantly deviates from that in shRNA dataset via a Fisher’s exact test on the contingency table. If each of the values in the contingency table was greater than 30, we used the chi-squared approximation of the Fisher’s exact test. We further filtered out any candidate CDE genes whose copy number was also significantly associated with the given mutation to control for potentially spurious associations coming from copy number (we removed genes showing significant association (FDR<0.1)). We performed this procedure for all 61 candidate genes one by one and selected those with FDR corrected Fisher’s exact test <0.1. We further filtered out the candidate master regulators whose mutation profile is correlated with *p53* mutation profile via a pairwise Fisher test of independence (FDR<0.1). We finally report the master regulators that have substantial number of CDE+ genes (N>300).

### Pathway enrichment analysis of CDE+/CDE− genes

We analyzed the CDE+/CDE− genes of each of the master CDE regulators for their pathway enrichment with the pathway annotations from the Reactome database^29^ in two different ways. First, we tested for significant overlap between our CDE genes with each of the pathways with hypergeometric test (FDR<0.1). Second, we ranked all the genes in the CRISPR-KO screen by the differences in their median post-KO cell viability values in mutant vs WT cells, and the standard GSEA method^30^ was employed to test whether the genes of each Reactome pathway have significantly higher or lower ranks vs the rest of the genes (FDR<0.1). We repeated the GSEA analysis with the genes ranked by differential post-KD cell viability in the shRNA screen, and only reported significant pathways specific to CRISPR but not shRNA screen. We confirmed that for *p53*, the GSEA method was able to recover the top significant pathways identified by the hypergeometric test (e.g. those in **Figure 1e**), although extra significant pathways were identified (**Table S3**). For *p53* and *KRAS* CDE− genes respectively, the enriched pathways were clustered based on the Jaccard index and the number of overlapping genes with Enrichment Map^31^, and the largest clusters were visualized as network diagrams with Cytoscape^32^.

To study the potential enrichment of CDE genes in common fragile sites (CFSs), we obtained the chromosomal band locations of CFSs from^14^, and we defined the CFS gene set as the set of all genes located within these chromosomal bands (obtained from Biomart^33^). We tested for a significant overlap between our CDE genes and the CFS gene set with hypergeometric test, and also confirmed the lack of significant overlap with the corresponding shRNA-DE genesets. Similarly for the common highly accessible chromatin (HAC) regions, we obtained a list of these regions defined by a consensus of DNAsel and FAIRE across seven different cancer cell lines from a previous study^34^. Next, we identified sgRNAs which are expected to target such HAC regions (see the *Calculating off-target scores* section below) and ranked genes based on the number of targeting such sgRNAs. Taking the top genes equal to the number of *p53* CDE+ genes, we computed the enrichment for *p53* CDE+ genes via a hypergeometric test.

### Testing the clinical relevance of copy number alterations of CDE genes in p53 mutated vs WT tumors

Given that the CRISPR-KOs of CDE+ genes preferentially reduce the viability of *p53* WT cells, we hypothesized that copy number alterations in CDE+ genes will analogously reduce the fitness of *p53* WT tumors. To test this hypothesis, we downloaded the somatic copy number alteration (SCNA) and patient survival data of 7.547 samples in 26 tumor types from cancer genome atlas (TCGA)^20^ from UCSC Xena browser (https://xenabrowser.net/). We confirmed that in these tumor types *p53* is mutated in more than 5% of the samples.

Our first goal was to study if the SCNA (both amplifications and deletions) in CDE+ genes is significantly lower specifically in *p53* WT tumors compared to that of control genes, and not in *p53* mutant tumors. As the control for our analysis, we used genes whose essentiality is not associated with *p53* mutational status. To this end, we computed the copy number alterations (genomic instability (GI)) of a given geneset, which aims to quantify the relative amplification or deletion of genes in a tumor based on SCNA. Given *s*_*i*_, the absolute of log ratio of SCNA of gene *i* in a sample relative to normal control, GI of the sample is given as in^35^:

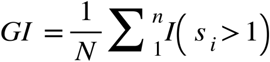

where I is the indicator function. We then checked whether the GI of CDE+ geneset is significantly lower than that of control non-CDE genes in *p53* WT but not in *p53* mutant tumors (using a Wilcoxon rank sum test).

Our second goal was to study if the magnitude of these SCNA levels of CDE+ genes is associated with *enhanced* patient survival, specifically in *p53* WT tumors, as this would further testify that such amplification/deletion events reduce the fitness of these tumors. To this end, we used the following Cox proportional hazard model to identify the genes whose genomic alteration is associated with patient survival specifically in *p53* WT tumor, while controlling for various confounding factors including the effect of cancer types, genome-wide genomic instability, sex, age, and race, tumor purity. We considered both amplification and deletion by taking the absolute value of SCNA levels,

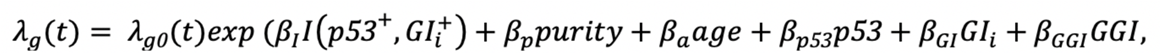

where *g* is an indicator variable over all possible combinations of patients’ stratifications based on cancer-type, race, and sex. *λ*_*g*_*(t)* is the hazard function (defined as the risk of death of patients per unit time) and *λ*_*g0*_*(t)* is the baseline-hazard function at time *t* of the *g*^*th*^ stratification. The model contains six covariates: (i) *I(p53*^*+*^,*GI*^*+*^), an indicator variable that has value 1 if *p53* is not mutated and the absolute value of SCNA level of the given gene *i* is greater than 50-percentile across all TCGA samples in the patient’s tumor, and otherwise 0, (ii) purity, denoting tumor purity^36^, (iii) age, denoting patient’s age, (iv) *p53*, denoting the *p53* mutational status of the patient, (v) GI_*i*_, the absolute value of SCNA levels of the given gene *i*, and (vi) GGI, quantifying the genome-wide genomic instability (GGI) of the sample, as computed above. We tested the enrichment of CDE+ genes among the genes whose absolute SCNA levels are significantly associated with better patient survival specifically in *p53* WT and not *p53*-mutated tumors using a hypergeometric test.

### Constructs and stable cell lines

*p53* R248Q was PCR amplified from a bacterial expression plasmid (kind gift of Dr. Shannon Laubert, UCSD) and *KRAS*G12D the pBabe-*KRAS*G12D plasmid (Addgene plasmid 58902, from Dr. Channing Der) using the Kappa Hi-fidelity DNA polymerase (Kappa Biosystems). These PCR amplicons were separately cloned into the MSCV-IRES-tdTomato (pMIT) vector (a kind gift from Dr. Hasan Jumaa, Ulm) using Gibson Assembly. We first generated high-efficiency Cas9-editing MOLM13 leukemia cells by transducing these cells with the pLenti-Cas9-blasticidin construct (Adggene plasmid 52962 – from Dr. Feng Zhang) and selecting stable clones using flow-sorting. Clones were then tested for editing efficiency by performing TIDE analysis^37^. These MOLM13-Cas9 cells were then transduced retrovirally with the pMIT-*p53*R248Q or pMIT-*KRAS*G12D mutants and sorted for tdTomato using flow-cytometry to generate isogenic mutant MOLM13-Cas9 cell lines.

### Generation of pooled sgRNA libraries

For pooled library cloning, 10 sgRNAs per gene were designed using the gene perturbation platform (https://portals.broadinstitute.org/gpp/public/analysis-tools/sgrna-design) Genetic Perturbation Platform. Guides targeting *p53* CDE+ and CDE− genes were synthesized as pools using array-based synthesis and cloned in the Lentiguide puro vector (Addgene plasmid 52963 – kind gift from Dr. Feng Zhang) using Golden Gate Assembly. A similar approach was used for the *KRAS* CDE libraries.

### Pooled sgRNA library screen

30 million MOLM13-Cas9 cells or their isogenic MOLM13-*p53* or *KRAS* mutant counterparts were transduced with the pooled CDE library virus in RPMI medium supplemented with 10% fetal bovine serum, antibiotics and 8 μg/ml polybrene. The medium was changed 24 hours after transduction to remove the polybrene and cells were plated in fresh culture medium. 48 hours after transduction, puromycin was added at a concentration of 1 μg/ml to select for cells transduced with the sgRNA library. Puromycin was removed after 72 hours and then cells were cultured for up to 30 days. 7 days after transduction, approximately 4 million cells were collected, and genomic DNA was prepared for the time zero (T0) measurement and also from time 30 (T30). Genomic DNA from these cells was used for PCR amplification of sgRNAs and sequenced using a MiSeq system (Illumina). Fold depletion or enrichment of sgRNAs from the NGS data was calculated using PinAplPy software^38^.

### CDE+/− genes identified in isogenic experiments

From the read counts per million for each sgRNA at Day 0 and Day 30 from the above pooled CRISPR screens across two replicates, we removed all the sgRNAs with read count < 20 at Day 0. We calculated an average fold change (FC) of reads from Day 0 to Day 30. For each sgRNA, we calculated this FC-rank difference in *p53* WT vs mutant in both CRISPR-KO and CRISPRi screens. For consistent comparison with AVANA, we only considered sgRNAs used in both libraries. The top and bottom genes are *differentially essential (DE)* from each screen. Taking the top ranked genes based on the difference of this score in two screens, we identify the CDE+ and CDE− genes.

### CRISPR Competition experiments

sgRNAs were cloned using standard cloning protocols and lentiviral supernatants were made from these sgRNAs in the 96-well arrayed format. 100,000 MOLM13 cells or tdTomato-positive isogenic mutants were plated in a 96 well pate and transduced with the sgRNA viral supernatants by spinfection with polybrene-supplemented medium. After selection of sgRNA transduced cells with puromycin for 48 hours, sgRNA transduced MOLM13 cells or mutants were mixed together in a ratio of 95:5 respectively, and the percentage of *p53* wildtype or *p53* mutant cells was monitored progressively up to 25 days using high-throughput flow-cytometry as described previously^39^.

## Supporting information

SuppMaterials

SuppTable1

SuppTable2

SuppTable3

SuppTable4

SuppTable5

SuppTable6

## Data and Code availability Statement

We have provided the scripts and data from both previously published and in-house screens, in their raw and processed form to reproduce each step of results and plots in a GitHub repository which can be accessed here: https://github.com/ruppinlab/crispr_risk.git

## Acknowledgements

We acknowledge and thank the National Cancer Institute for providing financial and infrastructural support. We thank Curtis Harris, Andre Nussenzweig, Sridhar Hannenhalli and the members of Cancer Data Science Lab for insightful feedback. This research was supported in part by the Intramural Research Program of the National Institutes of Health, NCI. S.S and K.C. are supported by the NCI-UMD Partnership for Integrative Cancer Research Program. AJD would like to acknowledge the support of the National Cancer Institute of the National Institutes of Health under Award Number P30 CA030199, the Rally Foundation for Childhood Cancer Research and Luke Tatsu Johnson Foundation under Award Number 19YIN45, an Emerging Scientist Award from the Children’s Cancer Research Fund, and the V Foundation for Cancer Research (TVF) under Award Number DVP2019-015.

## Notes

#### Summary of Updates

We validated our previous findings in a genome-wide manner by analyzing independent CRISPR screens and performed a new set of pooled and arrayed CRISPR screens to evaluate the competition between CRISPR-edited isogenic p53 WT and mutant cell lines.

https://github.com/ruppinlab/crispr_risk.git

