## Supplementary material for "Integrated computational and experimental identification of *p53, KRAS* and *VHL* mutant selection associated with CRISPR-Cas9 editing": SuppMaterials

Supplementary Notes

*1. Enrichment of p53 CDE+/- genes in the DE+/- genes from Haapaniemi et al.^3^*

To examine whether the *p53* CDE+/- genes can show similar effects during CRISPR gene editing in primary cells, we analyzed the original genome-wide screens in *p53* isogenic non-transformed RPE1 cells from Haapaniemi *et al.^3^*. We first observed that our knockout of our CDE+ genes identified from *AVANA* screens are more essential in *p53* WT cell than Mutant cells (P<0.02). Next, we ranked all the genes from the RPE1 cell screen by the difference of essentiality in WT vs isogenic *p53* mutant cells, and took the top *p* genes (i.e. that are more essential in WT, termed DE+ genes) and the bottom *q* genes (i.e. they are more essential in the mutant cells, termed DE- genes), where *p* and *q* are CDE+ and CDE- geneset sizes, respectively. We then tested for the significance of overlap between the CDE+ and DE+ genes with a hypergeometric test (similar for CDE- and DE- genes). To test this overlap enrichment robustly, we also used top *X* CDE+/- genes and top *Y* DE+/- genes, with *X* and *Y* ranging from 50 to the total number of CDE+/- genes with an interval of 50.

*2. Enrichment of KRAS CDE+/- genes in the publicly available KRAS isogenic essentiality screens*

To evaluate the relevance of our large-scale analysis in a single isogenic cell lines, we analyzed the CRISPR-KO and shRNA-KD screen performed in *KRAS* mutated isogenic DLD1 cell line^4^. In both the screens, we identified the CDE+ genes which are differentially more essential genes in WT vs Mut in CRISPR screens but not shRNA and similarly, we identified CDE- genes. For these CDE+ & CDE- (*Martin-CDE*) genesets, we tested their overlap enrichment with previously identified CDE+ and CDE- (*AVANA-CDE*) from pooled *AVANA* screens, respectively.

*3. Off-Target DNA damage effect on cell fitness is dependent on master CDE regulators*

Based on the notion that the master CDE regulators (MRs) could regulate the DNA damage response induced by Cas9, we hypothesize that the level of sgRNA off-target effect is associated with the extent of DNA damage response and consequently cell viability after gene knockout (KO), and this would be dependent on the mutation status of the MRs. Specifically, the level of sgRNA off-target effect will be negatively correlated with post-KO cell viability, but preferentially only in the MR-wildtype cells. Contribution of off-target to cell fitness is a combined effect of DNA damage and silencing of off-target genes. To avoid confounding by the latter, we considered only potential off-target hits in the non-coding regions and calculated an off-target score (0 to 1, where 0 represents no off-target hits) for each sgRNA. Specifically, for each sgRNA sequence, we calculated the genome-wide off-target score and list of potential off-target sites using CRISPRseek^1^. Top 100 potential off-target sites are taken into consideration to compute an overall cutting frequency determination off-target score^2^. We observed a significantly stronger positive correlation between gene essentiality and off-target score in MR-wildtype vs mutant cell lines (P<0.01 for *p53*, P<1.3E-04 for *VHL* and P<0.01 for *KRAS*). This difference in correlation strength further increases if we only consider top genes ranked by off-target score inducing a higher extent of DNA damage. These results suggest that similar to *p53*, the CDE effects of *VHL* and *KRAS* are also likely mediated by their potential role in DNA damage response.

Based on the above computed off-target score, we took *x* top ranked genes with highest off-target score and tested for their enrichment for the *CDE+* genes for each MR, where the value of *x* is the number of the respective *CDE+* genes. The fraction of CDE+ genes that are also among the top *x* genes with the highest off-target scores (summarized over all three MRs) was used as a measure of the accountability of CDE+ genes by sgRNA off-target effects.

List of Supplementary Tables

**Table S1**. *List of cell lines and genes present in both AVANA and Achilles, which has been used in this study.*

**Table S2**. *Differential essentiality analysis in p53 loss of function vs wild-type cell lines and the pathway analysis for the resulting CDE+/- genes*

**Table S3**. *CDE+ and CDE- genes and their enriched pathways*

**Table S4**. *Chromosomal common fragile sites (CFSs) enrichment analysis*

**Table S5**. *Fisher test significance of candidates from Vogelstein et al. which are mutated in at least 10 samples*

**Table S6**. *List of overlapping CDE+ genes across KRAS, VHL and TP53*

Supplementary Figures

| **a**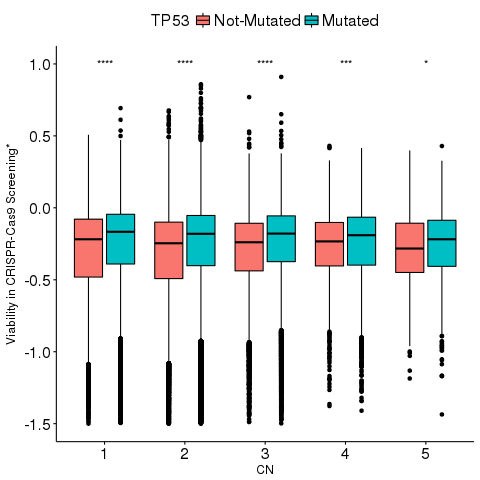 | **b**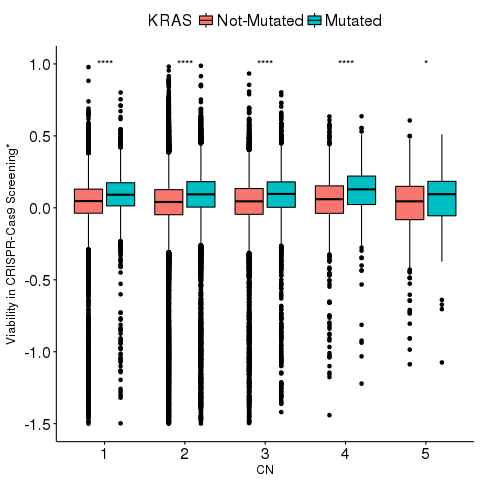 |
| --- | --- |

**Figure S1**. The effect of master regulators are independent of gene copy number.The x-axis shows the copy number and y-axis shows the cell viability after CRISPR-KO of each gene. Red bars denote the cell viability in the cell lines where the master regulator is WT, and green bars denote that where the master regulator is mutated for (a) *TP53* and (b) *KRAS*. CN=1 denotes cases with copy number less than or equal to 1 and CN=5 denotes cases with copy number greater than or equal to 5. The number of stars at the top of boxplot represents the significance of the difference (one-sided Wilcoxon rank sum test). *P<0.05, **P<0.01, ***P<0.001, ****P<0.0001.

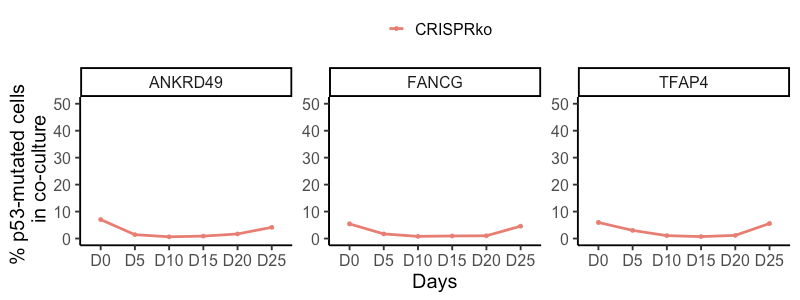

**Figure S2: Three CDE+ genes out of five showing** **no selection for *p53* mutant cells under CRISPR-Cas9 knockout of in a co-culture of *p53* WT/mutant cells.** In the competition assay, where isogenic *p53* WT/mutant MOLM13 cell lines were mixed with a ratio of 5:95 and top *p53* CDE+ genes were knocked out by CRISPR-Cas9. Change of ratio of *p53* mutant and WT cells (Y-axis, % *p53* Mutant/WT Cells) with time (number of days on the X-axis), where days are color coded. The leftmost panel (NTC) shows the corresponding result in the control experiment using non-targeting sgRNAs. Here, the y-axis scale is kept consistent to the main text.
